# GxP Single-cell RNA-seq and Spatial Transcriptomics end-to-end pipeline for clinical research

**DOI:** 10.64898/2026.01.23.701261

**Authors:** Amaya Zaratiegui, Timothy Burfield, Helle Rus Povlsen, Martín Emilio García Solá, Adrian Czaban, Keng Soh, Vivek Das

**Author notes:** Co-first author with equal contribution (Amaya Zaratiegui, Timothy Burfield).

## Abstract

Single-cell/nucleus RNA-sequencing and Spatial Transcriptomics are powerful tools for investigating cellular heterogeneity and tissue architecture that have deepened our disease understanding. Their broader adoption in clinical and regulated settings, however, is hindered by challenges related to data integrity, regulatory compliance, reproducibility, and scalability. To address this gap, we developed NNclinSSOAP (Novo Nordisk Clinical Single-cell Spatial Omics Analytical Pipeline) - a modular, GxP-ready end-to-end computational pipeline, that combines established single-cell workflows with a new Nextflow pipeline for Spatial Transcriptomics. NNclinSSOAP transforms RNA sequencing and Xenium spatial data into integrated, annotated single-cell objects and spatially resolved tissue maps. Designed to support mechanistic studies and clinical endpoint generation, it enables traceable and reproducible processing of large-scale datasets, scalable for both local and HPC environments. Here, we provide a step-by-step guide for using NNclinSSOAP. All code and data are publicly available. Using a standard laptop, the pipeline can be executed within 1.5 hours.

## Introduction

Single-cell and single-nuclei sequencing (sc/snRNA-seq) has revolutionized biological research by enabling the study of gene expression at the resolution of individual cells and nuclei. The technology allows for the identification of cell types, the characterization of cellular heterogeneity within tissues, underlying cell-cell communication and the understanding of complex biological processes at an unprecedented level of detail^1^. This granularity can uncover rare cell populations, specific gene expression profiles, and dynamic changes over time^2^. However, it inherently lacks spatial ^3^ information, which is critical for understanding the tissue architecture and the local microenvironment in which the cells reside^4^. Conversely, Spatial Transcriptomics retains high-resolution spatial information but often lacks the single-cell resolution necessary to discern the complexity of cellular interactions and the interplay of different cell types within tissues^5^.

The multi-modal omics approach of combining scRNA-seq and Spatial Transcriptomics addresses their respective limitations and provides a more comprehensive overview of cellular mechanisms in tissue contexts. This enables cellular identity and spatial organisation to be established concurrently, which can aid in revealing the mechanisms of action of drugs and understanding disease pathology^6–8^.

The use of these technologies, and the large data handling requirements that come with them, pose challenges for use in clinical settings. Ideally, such data needs to be stored and handled in a Good Experimental Practice (GxP) compliant way and compute resources need to be large enough to handle the data volumes. With this comes requirements on reproducibility of results and data provenance, leading to the development of modular scientific pipeline management and compliance tools such as Nextflow^9,10^, nf-core^11^ and BioCompute Objects (BCOs)^12,13^.

This paper addresses these critical challenges, describing the implementation and usage instructions for NNclinSSOAP ‐ a GxP-ready pipeline for running sc/snRNA-seq and Spatial Transcriptomics together. Our approach ensures data integrity and facilitates the translation of findings into robust and reliable conclusions for downstream applications, contributing to the standardization and quality control necessary for wider application of sc/snRNA-seq and Spatial Transcriptomics in regulated environments and clinical trial settings.

## Pipeline overview

The development of this paper was motivated by the need for a combined sc/snRNA-seq and Spatial Transcriptomics workflow in a clinical research and trial endpoint analysis context. While single-cell and Spatial Transcriptomics technologies have become widely used for tissue and disease understanding for hypothesis generation, their application in regulated environments such as clinical trials presents specific challenges. These include requirements for data integrity, traceability and reproducibility to comply with GxP standards. Existing workflows, although effective in research settings, often do not address these requirements in a systematic way.

To meet these needs, we designed an end-to-end pipeline that integrates established methods and best practices from the broader community, with adaptations that ensure GxP compliance and scalability. It was developed to ingest, process and analyse sc/snRNA-seq and spatial data to generate clinical endpoints and run initial analysis, while providing a transparent and auditable framework suitable for clinical studies. Due to the number of samples set to be collected from multiple omics modalities, it was clear that the workflow would need to handle large data volumes and support execution in a high-performance computing (HPC) environment. To address these points, we adopted a modular, containerized architecture using AWS, and our own in-house R environment.

To ensure data traceability and security, all data storage and access were managed using a compliant Amazon Web Services (AWS) Storage Service. Input and output data were handled in separate storage locations, with strict read and write access privileges limited to authorised users, thereby ensuring full adherence to regulatory requirements and maintaining data integrity. For data processing, analyses were performed within a dedicated, GxP-validated compute environment developed in-house as part of the Novo Nordisk secure analytics platform. This environment operates within a cloud-based infrastructure to provide both scalability and regulatory compliance throughout the data lifecycle.

Nextflow, an open-source workflow management system^10^, which enables reproducible and portable execution of complex bioinformatics analyses was used to process both sc/snRNA-seq and Spatial data. Each step was implemented as a separate module, allowing for both independent use and integration into larger workflows. A notable feature of the pipeline is the use of BioCompute Objects (BCOs), which provide a standardised, machine-readable record of computational analyses. By incorporating Nextflow’s BCO plugin, the workflow automatically records essential metadata from each run, supporting transparent reporting and communication with regulatory agencies and other stakeholders. This approach is intended to improve reproducibility and facilitate regulatory review.

The development process was iterative and collaborative, involving and building on cross-departmental expertise in computational biology, cloud engineering, and regulatory science. To address diverse data types and analytical requirements, NNclinSSOAP was designed in a modular fashion, that allows for straightforward adaptation to new datasets and evolving requirements, providing flexibility and scalability for future research. It is comprised of three main components, connected as illustrated in Figure 1 and described below:

**Figure 1.**
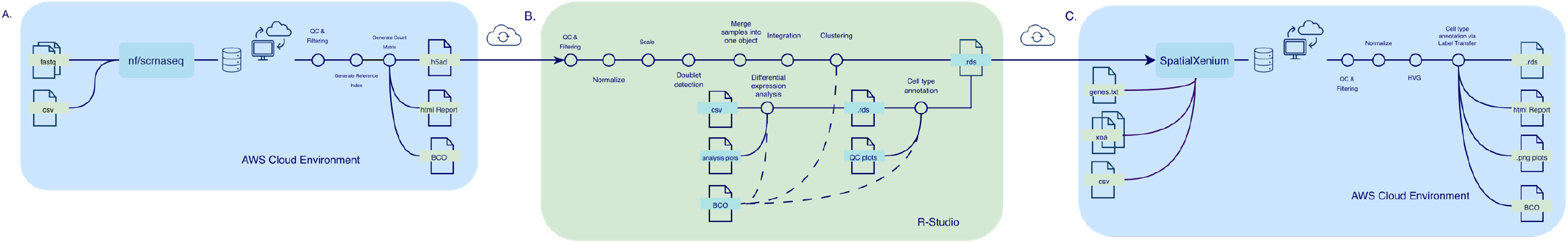
Overview of how the three main components of NNclinSSOAP are connected, with their respective inputs, processing steps, outputs, and compute environments. **a**, sc/snRNA-seq workflow. Processes fastq data, outputting, among others, h5ad files together with a multi-qc report and a BCO. The main processing steps include: Quality control, Generate reference index and Generate count matrix. **b**, R library/scripts workflow. Processes the h5ad file from a., outputting an annotated Seurat RDS object. The main processing steps include: Quality control, Doublet detection, Multiple sample integration, Seurat object generation, Differential expression analysis and Cell type annotation. **c**, SpatialXenium workflow. Processes raw Xenium data, outputting Spatial Transcriptomics analyses together with a report and a BCO, optionally taking the RDS object from the previous step to add cell type labels to the analyses. The main processing steps include: Quality control, Data normalization, Dimensionality reduction and clustering, Cell label transfer and Results visualisation.

a. sc/snRNA-seq workflow: Handles the preprocessing, alignment, and quantification of single-nucleus RNA-seq data, leveraging nf-core/scrnaseq^14^ for standardized and scalable data processing.
b. R library/scripts: Enables advanced post-processing, quality control, annotation and statistical analyses of the output from the sc/snRNA-seq pipelines, ensuring robust downstream interpretation.
c. SpatialXenium: A new Nextflow pipeline developed by us, designed to analyse multi-modal data from the Spatial Transcriptomics platform Xenium^15^, and optimised to run on HPCs ensuring efficient handling of large datasets. It ingests spatially-resolved gene expression data, performs quality control, normalization, clustering, and spatial niche construction. The pipeline outputs processed Seurat objects (RDS), cluster/niche compositions, top marker genes, and visualisations (PCA, UMAP, and spatial plots). Optionally, it performs label transfer, adding cell type annotation to the visualizations. The sub-modules that make up SpatialXenium are described in further detail in Box 1.

In practice, the Methods section in this paper demonstrates a test case that enables users to familiarise with NNclinSSOAP and convert raw sc/snRNA-seq and Spatial Transcriptomics data into spatially resolved tissue maps with gene expression and cell type annotations, through a small number of configuration changes and straightforward execution commands. The steps are divided into two procedures (Procedure 1 and 2) for sn/scRNA-seq and Spatial Transcriptomics respectively. These support local execution on the test datasets and outline the possibility of deployment on cloud infrastructure for larger datasets.

**Box 1**

**Brief introduction to the Spatial Transcriptomics processing steps (sub-modules)**

**1. SEURAT_XENIUM**

- Orchestrates reading/validating data, quality control, filtering, dimensionality reduction, clustering, and saving results.
- Can optionally produce a generically named output RDS if `--generic_name` is set to “TRUE”.

<u>Key Functions</u>

- *open_seurat()*: Loads Xenium data into Seurat, merging QC metadata.
- *plot_qc()*: Generates QC plots (e.g., read count distributions, gene feature counts).
- *filter_genes()*: Filters genes based on average quality value (QV) and read count thresholds.
- *run_seurat()*: Normalizes data, identifies variable features, runs PCA/UMAP/clustering.
- *get_cell_markers()*: Finds marker genes for each cluster.
- *plot_processed_data()*: Visualizes PCA, feature distributions, and UMAP.

**2. LABEL_TRANSFER**

- Reads the query Seurat RDS, reference RDS, and label column parameters.
- Calls *transfer_labels()* to annotate the Xenium data.
- Supports a `--future_mem_limit` parameter to set memory usage for parallel operations.
- Outputs labeled Seurat RDS and prediction files.

<u>Key Functions</u>

- *create_bygene_df()*: Summarizes gene counts across Seurat assays.
- *transfer_labels()*: Loads a reference RDS (single-cell data), ensuring matching label columns and overlapping features.
- Fits a linear model (query vs. reference gene counts), calculates residuals, and identifies anchors.
- Performs label transfer via *TransferData()*, generating predicted labels and probabilities.
- Filters low-probability cells (“Ambiguous”) and updates Seurat metadata.

**3. PLOT_FEATURE**

- Loads the Seurat object from RDS.
- Reads a text file of gene names.
- Produces and saves UMAP and spatial plots.
- If no gene list is provided, plotting is skipped.

<u>Key Functions</u>

- Generates `FeaturePlot` on UMAP for each specified gene.
- Creates `ImageFeaturePlot` to overlay gene expression on spatial coordinates.

### Applications

NNclinSSOAP was specifically developed for clinical trials incorporating transcriptomics as study endpoints, including sc/snRNA-seq and Spatial Xenium. As a pioneering framework in the pharmaceutical industry, particularly for cardiometabolic diseases, NNclinSSOAP has already been used in two mechanism-of-action trials. The first trial, focused on Alzheimer’s disease, incorporated scRNA-seq data as a primary endpoint^16^, and therefore utilized only Procedure 1 which is tailored for single-cell transcriptomics. In contrast, the second trial, a clinical study in kidney disease^17,18^, employed both Procedure 1 and 2 as it included both snRNA-seq and Xenium Spatial Transcriptomics data.

By enabling rapid, traceable, and reproducible data processing, this workflow sets a new standard for transcriptomic analysis in clinical trials. Looking ahead, it is broadly applicable to trials in other therapeutic areas, provided that compatible input data is available. Thanks to Nextflow’s modular architecture, the workflow can be easily adapted to accommodate additional analytical modules in the future.

Notably, the SpatialXenium Nextflow pipeline developed as part of this paper is now publicly available and can be integrated into new or existing Nextflow workflows, independently of the broader workflow described in the paper.

### Comparison to other methods

There are numerous tools and packages available to facilitate sc/snRNA Transcriptomics data preprocessing^19^. For sc/snRNA-seq we used the established nf-core pipeline scrnaseq as part of Procedure 1. Beyond the initial preprocessing steps, downstream processing and analysis of sc/snRNA-seq data commonly leverage widely used packages such as Seurat^20^ that we use as part of our R library. Our SpatialXenium pipeline is wrapped around the Seurat suite relevant for Spatial Transcriptomics data QC, pre-processing and analytics. Six main features make NNclinSSOAP different to other methods:

1. Modularity. NNclinSSOAP not only makes it possible to run the underlying sc/snRNA-seq and SpatialXenium workflows separately, but the workflows themselves build on Nextflow modules. This simplifies both using the workflows and adapting them to future needs as well as making them scalable, a key factor in data heavy settings.
2. GxP-readiness. Using Nextflow for building our workflows enables NNclinSSOAP to be both scalable and reproducible. Combined with documentation of the workflows through BCOs adding data provenance to the workflows, NNclinSSOAP meets the GxP requirements of being both reproducible and traceable. When used for GxP purposes, it allows the user to connect to a compute environment of choice, without compromising reproducibility, enabling analysis of data volumes at a much greater scale than in the test case described in this paper.
3. The SpatialXenium pipeline. There is a publicly available pipeline, nf-core/spatialxe, currently under development. It is a flexible image- and coordinate-based re-analysis framework, focused on controlling and improving Xenium data quality through multiple segmentation and spatial-data methods. SpatialXenium differs from this, instead focusing on post-processing and analysis of Xenium data. It provides users a fully specified, protocol-locked and easily reproducible end-to-end pipeline, implementing Spatial Transcriptomics analysis from Seurat for Xenium data.
4. We provide an R library capable of processing outputs generated by nf-core/scrnaseq, supporting standard Seurat workflows and downstream annotation. These scripts offer enhanced flexibility, incorporating tools such as Harmony^21^ and RPCA^22^ for multi-sample integration. For cell-type annotation, automated approaches like Azimuth^23^ or SingleR^24^ are supported, and user-defined reference datasets can be added. For differential treatment effect analysis, we use a pseudobulk^25^ approach by aggregating counts per sample and cell type, followed by statistical assessment using edgeR^26^, which enables robust testing of group-level differences while accounting for biological variability. Alternatively, other statistical methods, can be seamlessly integrated as needed. An overview of this workflow is illustrated in.
5. By leveraging the whirl package^27^ to execute the R library, the pipeline produces a BCO along with multiple log files—one corresponding to each R script. The package offers flexibility, enabling users to modify certain domains within the BCO and to select specific scripts to run as needed.
6. The R environment required for running these procedures is provided as a Docker container, ensuring that all necessary packages and dependencies are included and eliminating compatibility issues for the user. This approach allows the entire environment to be reliably loaded and the workflow to be executed as intended. However, one limitation is that package versions within the container may become outdated as new releases are made. Users also have the flexibility to set up and use their own custom environments if preferred.

## Experimental design

### Expertise needed for implementation

For users following the procedures step-by-step, no expertise is needed and none of the topics below need to be considered. However, some familiarity with single-cell transcriptomics, relevant training in statistical analysis of high-dimensional omics data and running analytics scripts is advisable for a smooth experience and ability to understand the outputs scientifically.

If adapting the workflow outlined in this paper to another dataset than the one specified, carefully consider the points below as well as the information and instructions provided in the code repository *README.md* and.md files in */docs*. For this, a broad understanding of R, Nextflow, cloud computing, containerisation and single-cell transcriptomics is needed.

### Sampling strategy-dependent analysis

The test datasets used in Procedure 2 contain a single sample each, one with healthy kidney tissue and the other with cancerous kidney tissue. This case facilitates analysis from the perspective of seeing differences in cell types and gene expression. Depending on a dataset’s sampling strategy, experimental design and analysis plans as often pre-registered in a clinical trial setting, other relevant analyses and visualizations are possible e.g. mitochondrial genes, cell cycle genes, biological sex related genes, batch effect contribution from metadata, etc. For samples taken from the same subject at multiple timepoints (longitudinal data), it is possible to analyse changes over time. For samples taken at a single time point for multiple subjects (cross-sectional data) it is possible to analyse the homo/heterogeneity within a population. Combining these two strategies creates a clinical trial scenario where NNclinSSOAP can be used to analyse subjects’ tissue characteristics over time as a response to treatment, comparing to a placebo group or among groups at single timepoint for disease understanding.

### Optional settings and configurations

Beyond the default settings and basic configurations required to run the test case in the Methods section, NNclinSSOAP can be customised through various command line flags and configuration files.

For the sc/snRNA-seq workflow, users can customize the metadata and configuration files to use with their own data, according to the nf-core/scrnaseq documentation: https://nf-co.re/scrnaseq/4.0.0/docs/usage/. For improved version control and streamlined collaboration, it is highly recommended to maintain a dedicated Git repository containing all related metadata and configuration files.

Once the nf-core/scrnaseq pipeline has completed, NNclinSSOAP provides users with flexible tools to further analyze and explore their data. Specifically, three main R scripts are included for downstream analysis, each designed with high flexibility in both parameters and methods. To customize or modify the pipeline execution, users need to edit the associated configuration file. Box 2 provides an overview of the config file, detailing the available options and their usage, with explanations of key settings. These configurable options provide users extensive flexibility for preprocessing, integrating, and annotating single-cell data within the NNclinSSOAP framework. Note that the RPCA integration method is computationally intensive and may not be suitable for local execution.

To load data into the pipeline, users must modify the *configs/read_samples.txt* file, ensuring it accurately lists each sample along with its corresponding metadata. It is essential to keep the column names and their order unchanged, as part of the script commands are hardcoded to this structure; any deviation will prevent the scripts from running correctly. All columns are intended to populate the Seurat object’s metadata, but if your dataset lacks certain variables (arm and visit) populate them with an NA and scripts 1 and 2 will still function, allowing you to generate an integrated and annotated Seurat object. However, script 3—designed for differential gene expression analysis in clinical settings to assess treatment response—specifically calculates treatment effects by assuming the data is longitudinal, with two timepoints at minimum and two distinct treatment arms (e.g., drug A and placebo). For this script to work, your visit variable labels must be “bl” (baseline) and “eot” (end of treatment), and the treatment arms should be labeled “placebo” and “treatment”. If your data does not fit this structure, script 3 cannot be executed.

Since the pipeline is run using the whirl wrapper, you can also adjust the *whirl/_whirl_preprocessing.yml* and *whirl/_whirl_analysis.yml* config files to edit BCO domains or exclude certain scripts from execution as needed.

For SpatialXenium, the genes shown in the UMAP and spatial feature plots are read from *assets/genes.txt* in the Spatial Transcriptomics workflow. By default, this file contains the genes needed to reproduce our results on the test data. Users should adapt this file if using other datasets, to reflect their genes of interest. Multiple command line options control which cells and genes are included in the UMAP and spatial feature plots, and how strict the filtering is. These options are explained in more detail in Box 3.

**Box 2**

**Key Settings of preprocessing workflow config file: *configs/preprocessing_workflow.yml***

\# Preprocessing workflow

Input data: Users can select among cell/gene matrices, filtered H5 matrices, or both filtered and raw H5 matrices by toggling the corresponding options. By default, the filtered H5 matrix is used. If both raw and filtered H5 matrices are provided, the pipeline automatically detects and removes ambient RNA using SoupX^29^. For multi-modal experiments, set the relevant variable to true and specify the modality type (default: “Gene Expression”).

*matrix_barcode_feats: false*

*h5_filtered: true*

*h5_raw_filtered: false*

*modality: true*

*modality_type: “Gene Expression”*

\# Parameters

Empty droplet filtering: Removal of empty droplets is implemented using DropletUtils^30^ and can be set via the DoEmptyDrop parameter.

*DoEmptyDrop: true*

General parameters: Users may initialize the project name and define thresholds for minimum and maximum number of detected features per cell, minimum cell count, mitochondrial RNA content cutoff, and clustering resolution.

*FDR_cutoff_EmptyDrop: 0.01*

*SeuratProjectName: “Test”*

*nFeatures_max: 5000*

*nFeatures_min: 500*

*nCells_cutoff: 10*

*percentMT_cutoff: 10*

*cluster_resolution: 0.6*

Feature plot genes: Specific genes to visualize in feature plots can be listed under the genes_of_interest key

*genes_of_interest:*

*-MAP7*

*-NES*

Integration settings: Multi-sample integration can be enabled/disabled. Supported methods include “rpca” and “harmony”. If using RPCA, users can specify anchor features.

*DoIntegration: true*

*integration_method: “harmony”*

*rpca_anchors_features:*

*-2*

*-3*

\# Cell annotation

Users can choose between SingleR or Azimuth for cell annotation and supply the appropriate reference dataset. Both methods automatically add metadata to the object: SingleR populates main_labels and fine_labels, while Azimuth adds predicted.celltype.l1, predicted.celltype.l2 and predicted.celltypel3. *annotation_method: “Azimuth”*

*singleR_ref: “dice”*

*azimuth_ref: “pbmcref”*

\# Input/Output Paths

Paths for sample manifest files, quality control outputs, Seurat object storage, and annotation results are specified for streamlined organisation and reproducibility.

**Box 3**

**Command line options for SpatialXenium**

`--min_reads_per_cell`

Sets the minimum number of reads per cell required for the cell to be included in downstream analysis and plotting. Cells with fewer reads than this threshold are considered low-quality or poorly captured and are excluded from the UMAP and spatial feature plots. Increasing this value makes the analysis more stringent (keeps only well-covered cells), while decreasing it is more permissive (includes more low-depth cells).

`--max_reads_per_cell`

Sets the maximum number of reads per cell allowed for a cell to be kept in the analysis. Cells with more reads than this threshold are often potential doublets, multiplets or technical outliers and are filtered out. This helps avoid cells with abnormally high counts dominating the variance structure in UMAP and other analyses.

`--max_control_prop_per_cell`

Controls the maximum allowed proportion of control probes/codewords (e.g. negative controls, spike-in controls) per cell. If a cell has a higher fraction of control signal than this value, it is considered low quality or problematic and is removed. This ensures that most of the counts per cell correspond to biological genes of interest rather than control features.

`--n_highly_var_genes`

Sets the number of highly variable genes (HVGs) to be selected by the Seurat function `FindVariableFeatures().` These HVGs are typically used for dimensionality reduction (PCA, UMAP) because they capture most of the biologically relevant variation. A larger number of HVGs can capture more subtle structure but may add noise; a smaller number yields a more compact representation but might miss fine-grained differences.

`--min_quality_per_gene`

Defines the minimum average quality value (QV) required for a gene to be kept in the dataset. Genes whose average QV is below this threshold are filtered out as low-confidence or noisy measurements. This helps ensure that only genes with sufficiently reliable signal contribute to the UMAP and spatial feature plots.

`--min_celltype_probability`

This parameter is used when label transfer is executed. It controls the minimum assignment probability required for a cell to be labelled as a specific cell type. If the predicted probability for the best-matching cell type is below this threshold, the cell is assigned the label “Ambiguous” instead of a specific type. Lowering this value will assign more cells to named types (more aggressive labelling), while increasing it will produce more conservative assignments (fewer cells confidently labelled, more “Ambiguous”).

### Seurat Spatial Transcriptomics visualisation options

The tissue visualisations overlayed with gene expression and cell types generated from running SpatialXenium utilise the Seurat v5 package’s functions `ImageDimPlot()`and `ImageFeaturePlot()`. These offer a wide range of options beyond what is covered by default in SpatialXenium, enabling extensive control over plot aesthetics and content, including colour scales, transparency, cell border properties, molecule overlays, co-expression blending for pairs of genes and grouping cell types. For advanced users wanting to tailor SpatialXenium visualisations to their specific biological questions, `ImageDimPlot()` is called from *label_transfers.r* and `ImageFeaturePlot()` from *plot_features.r*. We strongly recommend familiarity with R, Seurat and Nextflow for doing this, as making changes may result in errors downstream.

A vignette outlining the visualisation possibilities can be accessed here: https://satijalab.org/seurat/articles/seurat5_spatial_vignette_2.html

### Compute environment

NNclinSSOAP was optimised to run on high-performance computing environments. For running the minimal test case described in this paper the hardware requirements are basic enough to use a laptop. However, this is unlikely to be the case if attempting to analyse larger datasets, such as the benchmark datasets listed in the Resource availability section. Benchmarking results from running the SpatialXenium workflow on these in our cloud environment are listed in Table 1.

**Table 1.** Benchmark results for the SpatialXenium workflow. Computational requirements and duration for running the benchmark datasets in our cloud environment. As the Xenium input size increases, the RAM required for analysis quickly exceeds what is commonly available in a local compute environment.

| Dataset | Xenium Input Size | Output Size | Duration | Max RAM required |
| --- | --- | --- | --- | --- |
| Kidney non-diseased | 9.4 GB | 695 MB | 13m 37s | 11.9 GB |
| Kidney cancer (PRCC) | 6.4 GB | 449 MB | 10m 17s | 10.2GB |
| Human Skin Melanoma | 12 GB | 2.3 GB | 16m 18s | 14.8 GB |
| Pancreatic Cancer | 7.8 GB | 3.1 GB | 25m 3s | 18.1 GB |
| Cervical Cancer 5k | 69 GB | 6.6 GB | 4h 38m 33s | 75.7 GB |

Memory and CPU allowances for different processes in the SpatialXenium pipeline can be customised in *conf/base.config*. The pipeline also allows usage of alternative containerisation platforms to Docker, such as Singularity, through the `-profile` Nextflow argument. In addition to this, Nextflow has options for connecting executors such as HPC/cloud computing services by editing *nextflow.config*, further explained here: https://www.nextflow.io/docs/latest/executor.html

## Materials and equipment setup

To meet all prerequisites for following the Methods section, ensure that all the requirements in this section are met, and any given instructions followed.

### Hardware requirements

- Any desktop workstation or laptop with an internet connection is sufficient. The procedures were tested on a MacBook Pro (MacOS Tahoe 26.1) with a 12-Core central processing unit (CPU) and 24 GB of random-access memory (RAM). For minimal performance, we recommend using 4 CPUs with at least 18 GB of RAM for analyses.

### Software requirements

- Operating system: Linux, Windows or MacOS
- IDE: An integrated development environment. We recommend Visual Studio Code which can be accessed at https://code.visualstudio.com/download.
- Docker desktop: A platform for docker container management, which can be accessed at https://docs.docker.com/desktop/.
- Nextflow: A workflow system for creating scalable, portable, and reproducible workflows, which can be accessed at https://www.nextflow.io/docs/latest/install.html. The version used here is 25.10.0.

### Installation of Docker desktop

Install Docker desktop from the official website https://www.docker.com/products/docker-desktop/. Once downloaded, open the application and go to Settings > Resources and change the CPU setting to a minimum of 4 and the memory setting to a minimum of 18GB.

### Installation of Nextflow

Install Nextflow by following the instructions on the official website https://www.nextflow.io/docs/latest/install.html. You can check if Nextflow is correctly downloaded and available by typing the following in your terminal: `nextflow info`

### Cloning the repository

Open a terminal and navigate to the directory you want to place the repository in, then run:

> git clone https://github.com/NovoNordisk-OpenSource/nnclinssoap.git

### Downloading the test data

Inside your IDE of choice, open the cloned repository. In the terminal, run the following:

`> bash./setup_test_data.sh`

You should see the download start in the terminal. Once finished (approximately 1h), the test data is found in the directories *scrnaseq/test_data* and *SpatialXenium/test_data*.

### Building the docker images

Start by making sure Docker desktop is open. Then, inside you IDE of choice, open the cloned repository. In the terminal, run the following:

`> cd docker`

`> bash./build_all_images.sh -v docker`

Build progress is shown in the terminal and Docker desktop. Building all images takes around 3-6h.

## Methods

This section describes how to use NNclinSSOAP with the provided test data, ensuring an easy-to-follow introduction to usage. For users who wish to customise their analysis, use other data or run the pipeline in an HPC, the repository is open source and contains more detailed documentation.

We divide the methods in two procedures, which can be performed sequentially or independently. Procedure 1 describes the preparation and steps needed for running the sc/snRNA-seq workflow. Procedure 2 covers the preparation and steps needed for running the SpatialXenium workflow, with an option to incorporate the annotated Seurat object generated from Procedure 1 as input for cell label transfer.

If problems arise whilst following the procedure, please refer to the Troubleshooting section.

### Procedure 1: sc/snRNA-seq ~ 1h

#### Run nf-core/scrnaseq preprocessing (test execution) ~ 10 min

1. Open the nnclinssoap repository in your IDE.
2. In the terminal, run the following: `> nextflow run nf-core/scrnaseq -profile test,docker –outdir./results`
3. Once the pipeline has run, the output files will be placed under the output folder */results*. **Run the pipeline ~ 50 min** **Note:** Docker desktop must be open.
4. Run counts matrix processing into Seurat object. In the terminal, run the following: `> cd scrnaseq` `> bash./run_preprocessing_containerized.sh`
5. Run cell annotation and differential expression analysis. In the terminal, run the following:

`> bash./run_analysis_containerized.sh`

### Procedure 2: SpatialXenium ~ 15min

#### Set configurations ~ 3 min

1. Open the nnclinssoap repository in your IDE.
2. Open *SpatialXenium/conf/base.config* and check if any of `withLabel:process_single`, `withLabel:process_low`, `withLabel:process_medium` or `withLabel:process_high` have a cpu or memory setting exceeding your available resources and the settings in Docker Desktop. If so, change them to be within that range.
3. (Optional) Open *SpatialXenium/conf/test.config* and change `reference_rds` to the path of the annotated Seurat object created in Procedure 1, step 5 (should be *scrnaseq/output/seurat_object.rds)*. In the same file, change `single_cell_label_col` to `“predicted.celltype.l1”`. **Run the pipeline ~ 12 min** **Note:** Docker desktop must be open.
4. In the terminal, run the following: `> cd SpatialXenium` `> nextflow run main.nf -profile test,docker --outdir test_results`
5. Once the pipeline has run, the output files will be placed under the output folder */test_results*.

## Troubleshooting

For general troubleshooting there are shell scripts designed to validate that the pre-requisites for running NNclinSSOAP are in place, located in the directory */tests*. Run them from the terminal and read through the resulting logs. The scripts are described below:

- *test_containers.sh*: Container testing Suite. Tests all built containers for correct functionality.
- *spatialxenium_preflight_check.sh*: Pre-flight check for SpatialXenium pipeline testing. Verifies all dependencies and setup before running tests.

In case running these scripts does not indicate where your issue lies, refer to Table 2 for further troubleshooting related to the specific steps in the Methods section.

**Table 2.**
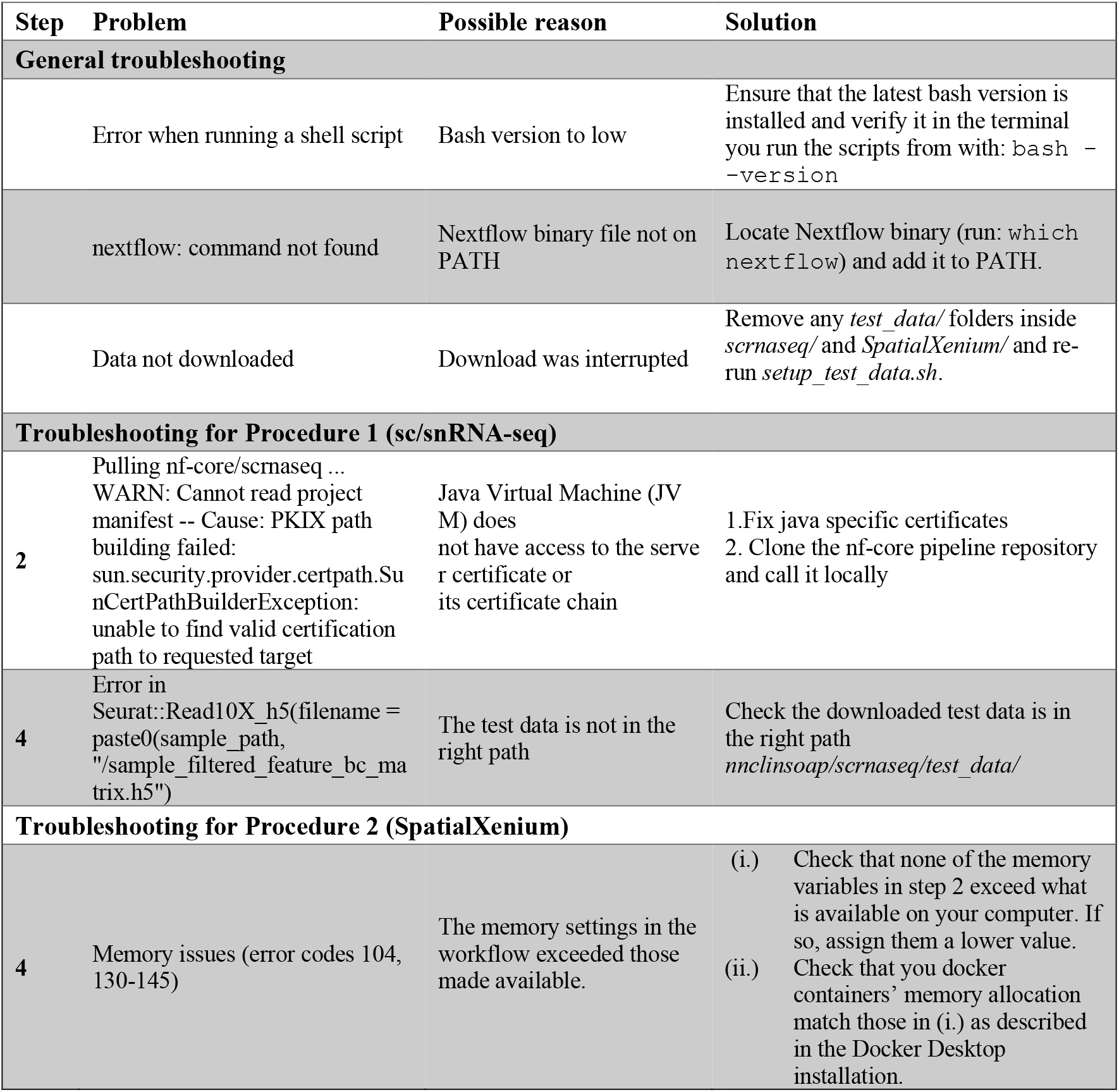
Step-specific troubleshooting. A collection of problems related to the different steps in the Methods section, with possible reasons and solutions.

## Expected results

The result of following both procedures is a transformation of raw sc/snRNA-seq data into an integrated and annotated Seurat object, and raw Xenium data into a spatially resolved cell object and map. This can aid in answering clinical questions, giving the user a deeper understanding of the analysed tissues both at a cellular expression and structural level, as is the case in the test data in this paper, highlighting differences between healthy and cancer kidney tissue.

Since reproducibility is at the core of this paper, the results produced by following both procedures are expected to be similar to those shown in this section. If the paper is followed using a personal dataset, the same types of outputs are expected but will likely differ significantly from those shown in this section. The entire workflow, including input, output, and parameters used is documented in a BioCompute Object. As BioCompute is an FDA-supported standard for regulatory submissions, it can be used during regulated activities and communication with the Agency.

### sc/snRNA-seq (Procedure 1)

Executing the sc/snRNA-seq workflow (Procedure 1, step 2) generates an nf-core results folder. The test dataset has been run with the cellranger aligner^28^ so the results are the standard cellranger outputs. The special addition is the BCO object that has been generated since the plugin is included when running the nextflow pipeline.

In the second part of the workflow (Procedure 1, step 4 & 5), each script produces a variety of plots and output files, which are organized into dedicated folders—see Box 4 for a summary of the results file structure.

- Quality Control (QC): QC plots document the standard Seurat workflow, starting from reading the counts matrix and progressing through to the creation of the final Seurat object (see Figure 2 a-d). A separate folder is generated for each sample’s QC outputs, while the integrated Seurat object is stored in the parent directory.
- Cell Annotation: This step generates visualizations such as UMAPs displaying annotated clusters and violin plot displaying the annotation scores, along with an updated Seurat object that contains cell type assignments (see Figure 2 e and f).
- Differential Expression analysis: For each cell type, the pipeline outputs a normalised counts CSV file, a CSV file with differential expression results, and a volcano and line plot summarizing key findings (see Figure 2 g and h).

**Figure 2.**
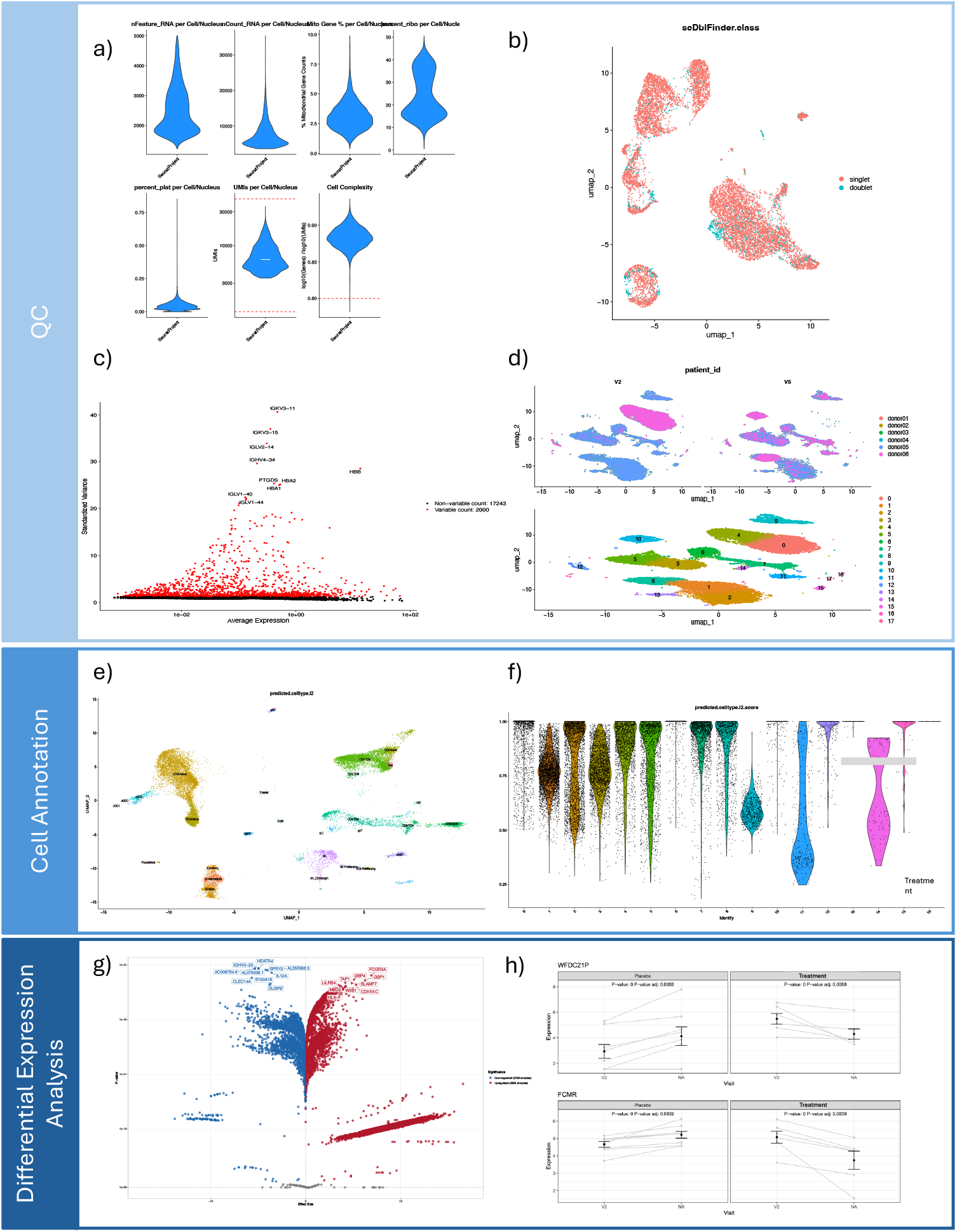
Summary of sc/snRNA-seq data anlysis workflow and results. **a**, QC violin plots displaying the distribution of nFeature_RNA, nCount_RNA, mitochondrial percentage, ribosomal percentage, platelet percentage, UMI counts, and cell complexity per cell. **b**, UMAP projection highlighting doublet detection results. **c**, Identification of top highly variable genes used for downstream analysis. **d**, Integration UMAP visualizations, stratified by patient ID (top) and by cluster (bottom). **e**, UMAP embedding with annotated cell types. **f**, Violin plots showing prediction scores for cell type assignment. **g**, Volcano plot displaying differential expression analysis results illustrating treatment effects. **h**, Line plot illustrating the averaged expression levels of the top differentially expressed genes across treatment groups.

### SpatialXenium (Procedure 2)

Executing the SpatialXenium workflow (Procedure 2, step 4) produces several key outputs that can be used for clinical interpretation of the tissues analysed, shown in Figure 3. By using Nextflow, reports covering errors, timing and resource usage have automatically been created. An example results file structure is shown in Box 5. To aid with technical and clinical interpretation, key outputs from Quality Control, gene expression and cell type analysis are described in further detail below:

**Figure 3.**
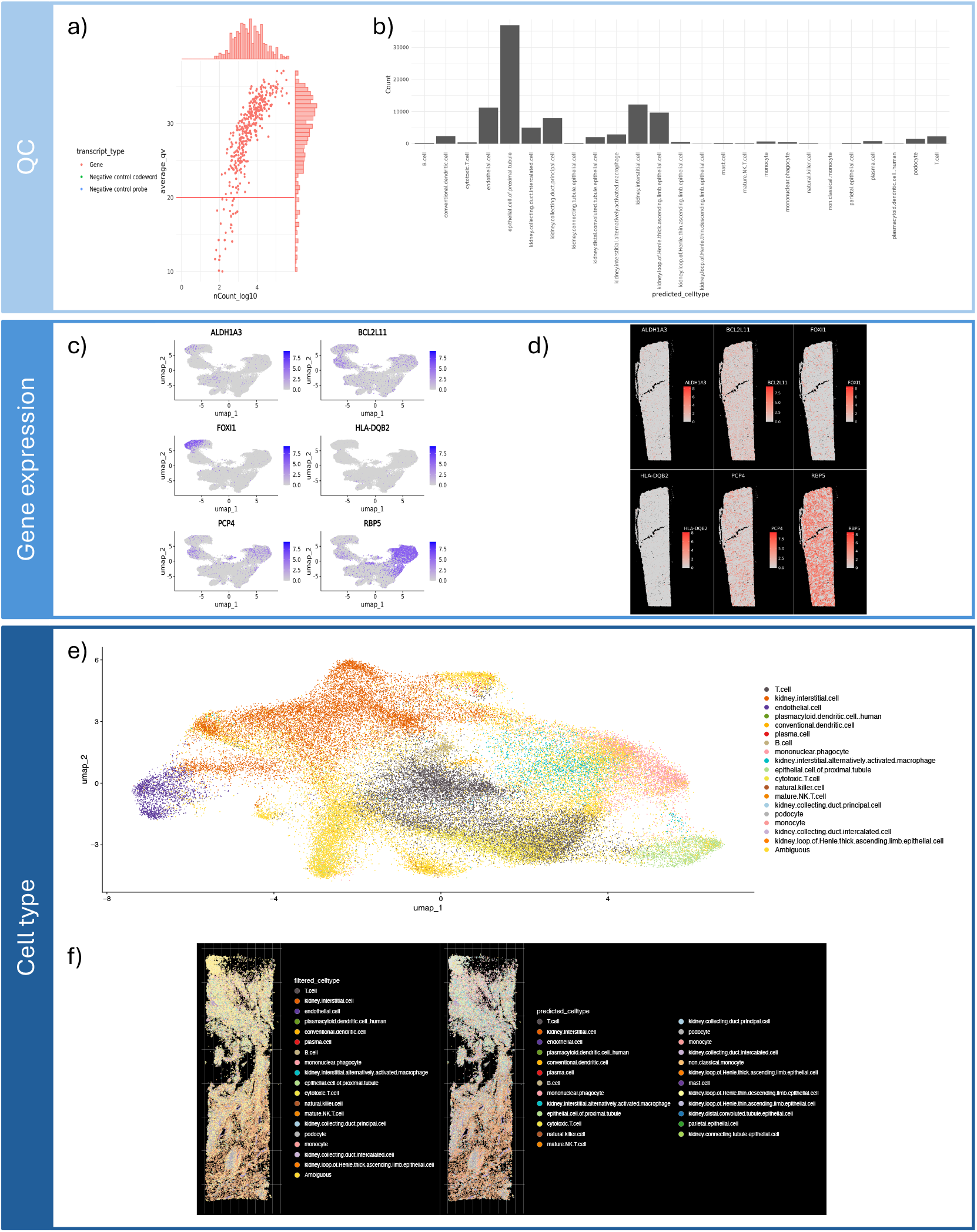
Visualizations generated by the Spatial Transcriptomics pipeline relating to Quality Control, Gene Expression and Label Transfer. **a**, scatter plot of the QV score vs the counts of all genes, with histograms demonstrating the distribution of genes in each axis. **b**, histogram showing the count of each predicted cell type. **c**, UMAPs highlighting the density of each gene (gradient of purple - blue) from genes.txt in the UMAP space containing all genes (grey area). **d**, Tissue maps highlighting the location and level of gene expression (gradient of red) for each gene in genes.txt. **e**, UMAP showing all the cell types, set to ‘Ambiguous’ below the probability threshold specified. **f**, Tissue maps with predicted cell types filtered by the specified probability threshold highlighted (left) and all predicted cell types (right).

- Quality Control (QC): The QC plots (see Figure 3 a and b) summarise how many genes and reads are captured per cell and how these values are distributed across the dataset. This allows users to identify low-quality cells, technical outliers or regions with suboptimal capture efficiency before drawing any biological or clinical conclusion, ensuring that downstream patterns (e.g. “immune poor” areas) are not artefacts of poor data quality.
- Gene Expression: The gene expression visualisations (see Figure 3 c and d) show how the selected markers from *assets/genes.txt* are distributed both in UMAP space and across the tissue. Clinically, these plots help confirm that canonical markers (for example tumour, immune or stromal genes) are expressed in the expected tissue compartments and spatial niches, supporting interpretations such as tumour–stroma interactions, immune infiltration patterns or localisation of specific disease-relevant pathways.
- Cell Type: The cell type plots (see Figure 3 e and f) resulting from the label transfer, display the predicted cell types in UMAP space projected back onto the tissue, with cells below the chosen probability threshold set to “Ambiguous”. This makes it clear which assignments are robust enough to support biological or clinical interpretation, which should be treated with caution. In a clinical context, these maps can be used to explore how specific cell types (e.g. cytotoxic T cells, fibroblasts, endothelial cells) are spatially organized within lesions, margins or normal tissue, and how this architecture might relate to disease severity or therapeutic response.

**Box 4**

**Example file structure from Procedure 1.5**

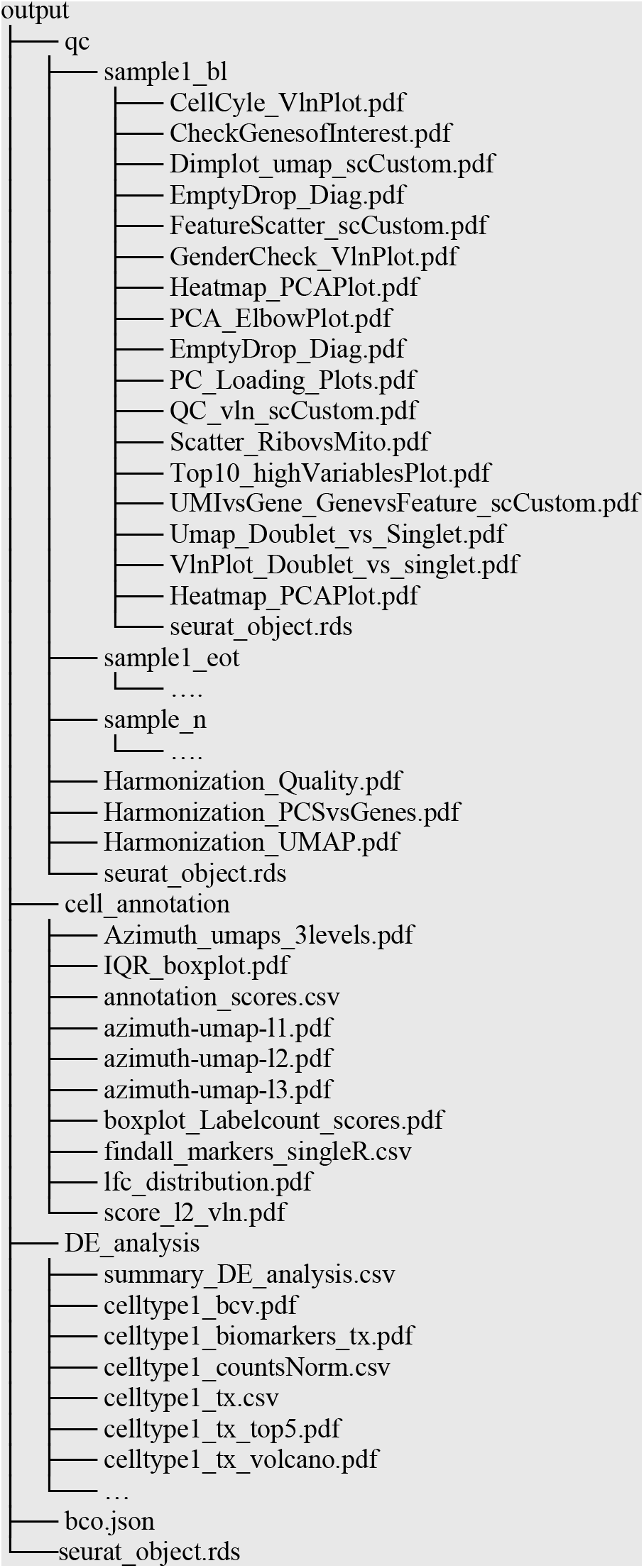

**Box 5**

**Example file structure. Full sample file structure only shown for sample 1 for simplicity**.

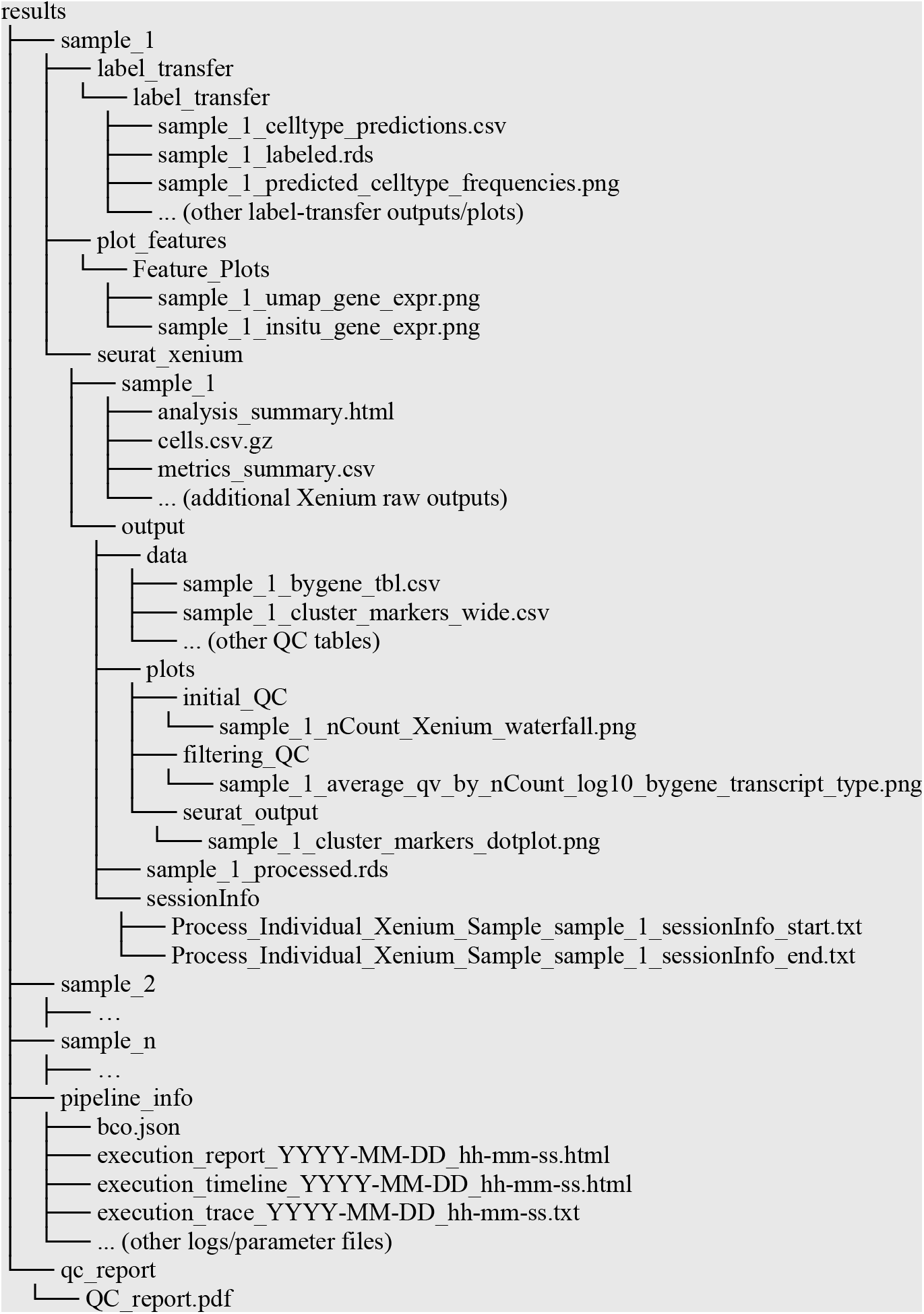

## Resource availability

### Lead contact

Requests for further information and resources should be directed to and will be fulfilled by the lead contact, Vivek Das.

### Data and code availability

#### Code

The NNclinSSOAP repository is available here: https://github.com/NovoNordisk-OpenSource/nnclinssoap.

#### Test data used in Methods section

All data used in the Methods section are available from the links listed below:

- scRNA-seq h5 matrixes: https://zenodo.org/records/18173543.
- Xenium Kidney (both non-diseased and cancer): https://www.10xgenomics.com/datasets/human-kidney-preview-data-xenium-human-multi-tissue-and-cancer-panel-1-standard.
- Human kidney cell annotation reference: https://zenodo.org/records/10694842.

#### Benchmark datasets (optional)

Datasets used for benchmarking purposes are available from the links listed below:

- Human Skin Melanoma: https://www.10xgenomics.com/datasets/human-skin-preview-data-xenium-human-skin-gene-expression-panel-1-standard.
- Pancreatic Cancer: https://www.10xgenomics.com/datasets/pancreatic-cancer-with-xenium-human-multi-tissue-and-cancer-panel-1-standard.
- Cervical Cancer 5k: https://www.10xgenomics.com/datasets/xenium-prime-ffpe-human-cervical-cancer.

#### Fastq dataset (optional)

Raw data for creating the scRNA-seq h5 matrixes using nf-core/scrnaseq is available from the links listed below:

- Universal 5’ dataset: https://www.10xgenomics.com/datasets/5-hashing-example-with-tabs-2-standard.

## Acknowledgement

VD conceived and conceptualized the study. AZ, TB, HP, MG built the analytical framework, generated and tested all codes related to execution of the software, pipeline, and setting up compute environment. AZ and TB ran all the analysis and generated all the figures. AC assisted around the regulatory compliance aspect. AZ and TB wrote the first draft with inputs from MG, AC, KS under the supervision of VD. All authors read, revised, validated and agreed to the content of the manuscript.

The authors extend their gratitude to Oskar Hint and Sebastian Dengler from Novo Nordisk A/S for their inputs related to AWS and compute environment.

